# Maximum Caliber can build and infer models of oscillation in a three-gene feedback network

**DOI:** 10.1101/444307

**Authors:** Taylor Firman, Anar Amgalan, Kingshuk Ghosh

## Abstract

Single-cell protein expression time trajectories provide rich temporal data quantifying cellular variability and its role in dictating fitness. However, theoretical models to analyze and fully extract information from these measurements remain limited for three reasons: i) gene expression profiles are noisy, rendering models of averages inapplicable, ii) experiments typically measure only a few protein species while leaving other molecular actors – necessary to build traditional bottom-up models – unnoticed, and iii) measured data is in fluorescence, not particle number. We have recently addressed these challenges in an alternate top-down approach using the principle of Maximum Caliber (MaxCal) to model genetic switches with one and two protein species. In the present work we address scalability and broader applicability of MaxCal by extending to a three-gene (*A*, *B*, *C*) feedback network that exhibits oscillation, commonly known as the repressilator. We test MaxCal’s inferential power by using synthetic data of noisy protein number time traces – serving as a proxy for experimental data – generated from a known underlying model. We notice that the minimal MaxCal model – accounting for production, degradation, and only one type of symmetric coupling between all three species – reasonably infers several underlying features of the circuit such as the effective production rate, degradation rate, frequency of oscillation, and protein number distribution. Next, we build models of higher complexity including different levels of coupling between *A*, *B*, and *C* and rigorously assess their relative performance. While the minimal model (with four parameters) performs remarkably well, we note that the most complex model (with six parameters) allowing all possible forms of crosstalk between *A*, *B*, and *C* slightly improves prediction of rates, but avoids ad-hoc assumption of all the other models. It is also the model of choice based on Bayesian Information Criteria. We further analyzed time trajectories in arbitrary fluorescence (using synthetic trajectories) to mimic realistic data. We conclude that even with a three-protein system including both fluorescence noise and intrinsic gene expression fluctuations, MaxCal can faithfully infer underlying details of the network, opening up future directions to model other network motifs with many species.

## Introduction

Advances in microfluidics and microscopy allows us to sensitively monitor gene expression profiles in live single-cells over unprecedentedly long times and at frequent intervals.^1–5^ These single-cell temporal measurements of gene expression are fundamentally different from transcriptomic and proteomic data that pool results from many cells and are not designed to preserve or track individual cells over long periods of time.^1^ Consequently, these measurements do not provide fluctuating time series data like single-cell measurements, nor do they allow us to get a glimpse of the underlying heterogeneity dictating fundamentals of biological regulation. ^6,7^ For example, fluctuations at the single-cell level can teach us how microbes and/or cancer cells evade drugs using noise and cell-cell variability.^8–12^

These measurements now demand new quantitative models to maximally harness the information hidden in long single-cell trajectories individually recorded for many cells. Traditional modeling schemes remain limited for two reasons. First, single-cell gene expression data is inherently noisy due to the small number of molecules involved. ^6, 13–25^ It follows that traditional models of averages based on mass-action type approaches ^26–31^ are not useful to explain the noisy data; stochastic models are inherently necessary. Next, stochastic models are typically bottom-up, requiring information about more species than experiments can measure. For example, these bottom-up models require information about mRNA, promoter-protein complexes, etc., while experiments usually only measure the expression level of key proteins, leaving other molecules unnoticed. The challenge of partial information gets even more demanding when different proteins interact with each other. In bottom-up models, feedback is often modeled by invoking auxiliary species like protein-protein complexes that are rarely observed in experiments.

We have recently introduced an alternate top-down approach based on the principle of Maximum Caliber (MaxCal) to analyze genetic circuits.^32–35^ MaxCal maximizes the path/trajectory entropy subject to minimal constraints to predict the trajectory probability distribution^33,36^ and circumvents both of the challenges mentioned above. First, MaxCal is built in the language of trajectories, thus it starts directly with the raw data. MaxCal generated path probabilities are used to maximize the likelihood of the experimentally observed trajectory. Thus, the joint MaxCal + maximum likelihood (ML) approach provides a much needed stochastic framework that harnesses the maximum amount of information from the single-cell trajectories recorded in experiments. Next, because of its top-down nature, it builds a minimal model without any ad-hoc assumptions. In our recent work, we have shown the success of this novel MaxCal + ML approach in describing two types of switch-like genetic networks: 1) a single gene auto-activating circuit,^34^ and 2) a two-gene mutually repressing genetic toggle switch.^35^ MaxCal has been successful in inferring underlying details of these circuits such as the effective production rate, degradation rate, and feedback that are not typically visible in experiments.

With the success of these studies, new questions emerge: How scalable is MaxCal, i.e. is MaxCal restricted to only one-gene and two-gene circuits? Much of biochemistry is complex, involving *cycles* of more than two states (as in molecular motors and pumps) or involving more than two proteins. Transcriptional regulation in gene expression networks entails multiple interconnected components and feedbacks.^37^ Can we use MaxCal to model such a scenario that involves networks with more than two genes? Furthermore, is the success of MaxCal limited to describing switch-like circuits where one gene toggles between the ‘ON’ (high expression) and ‘OFF’ (low expression) state? Can we describe circuits where gene expression levels vary in less abrupt manner? For example, can we describe oscillatory circuits, a basic feature of many natural circuits? Synthetic biologists are also designing new circuits beyond switch-like circuits to understand fundamentals of evolution and biological function.^38–44^ Motivated by these questions, we have chosen to analyze the repressilator circuit^45,46^ that arises from three-gene (*A*, *B*, *C*) circuits where gene *A* represses expression of gene *B*, *B* represses *C*, and *C* represses *A* (see Figure 1). This circuit is known to exhibit oscillation. We generated synthetic time series data for proteins *A*, *B*, and *C* from a known underlying model with nine species (see equation 1). We test MaxCal’s ability to infer network details by only providing time trace data for the three protein species. This exercise serves three purposes: i) it shows that MaxCal can model oscillatory circuits, beyond switches ii) it addresses the scalability of MaxCal by extending from two species to three species that interact with each other via a complex feedback network, and iii) it reiterates MaxCal’s ability to infer underlying circuit properties even when there is only partial data available. In this case, MaxCal uses information about the expression profile of three proteins while the underlying circuit has nine species.

**Figure 1:**
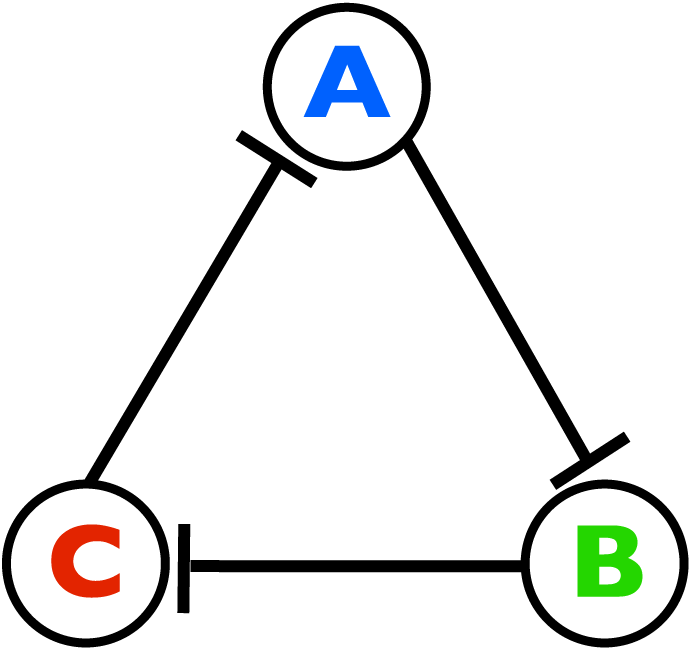
An oscillator circuit^45^ where the presence of protein *A* inhibits production of protein *B*, the presence of protein B inhibits production of protein *C*, and the presence of protein *C* inhibits production of protein *A*.

Next, we addressed another typical challenge associated with these measurements when input data is not given in discrete protein number, but in noisy fluorescence. Fluorescence per protein is not constant but a fluctuating quantity^47–49^ due to photobleaching, GFP maturation,^50,51^ and day-to-day variability.^52^ How do we analyze such noisy fluorescence data? The MaxCal + ML approach is particularly advantageous due to its reliance on raw trajectory. We have augmented the likelihood function by incorporating the fluorescence per protein distribution. As in our previous examples with genetic switches,^34,35^ we now demonstrate MaxCal + ML can reasonably infer details even when data is in florescence for all three proteins. Below, we first give the description of the repressilator circuit and the underlying model used to generate the synthetic data, followed by a description of our MaxCal model at different levels of complexity. Finally, in the results section, we present the inference power of these models and their relative comparison.

## Materials and Methods

### Model for repressilator circuit to generate synthetic data using Gillespie algorithm

To demonstrate MaxCal’s ability to model oscillatory circuits, we generated synthetic data of protein expression levels from a known model and benchmarked MaxCal’s predicted rates against these known values. Protein expression time trajectory data is generated by using a three-gene oscillatory circuit known as the repressilator.^45,46^ This circuit is explicitly defined by the following reaction scheme following the work of Loinger and Biham: ^46^

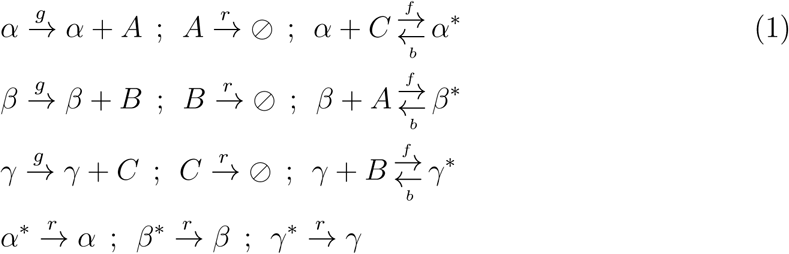

where generic proteins *A*, *B*, and *C* are created from their respective genes *α*, *β*, and *γ* at a rate of *g* and are degraded at a rate of *r*. The oscillatory behavior of the circuit is ensured by imposing the feedback scheme where *A* can bind and unbind the repressor site of *β*, *B* can bind and unbind the repressor site of *γ*, and *C* can bind and unbind the repressor site of *α* at rates of *f* and *b* respectively. Binding converts *α*, *β*, and *γ* into their deactivated states, *α^*^*, *β^*^*, and *γ^*^*, where they cannot create their respective proteins. In order to allow oscillation, proteins that are bound are also allowed to degrade at rate *r*. Rate values are chosen to reproduce oscillatory periods and heights that are representative of experiments while maintaining realistic protein synthesis and degradation rates (see Table 1). This circuit was simulated using a Gillespie algorithm^53^ to generate stochastic trajectories for expression of proteins *A*, *B*, and *C*. This serves as our input data to mimic experiment that will be used to benchmark MaxCal’s inference power. Protein numbers are sampled in increments of Δ*t* over the course of time length *T* to mimic experimentally realistic sampling rates.

**Table 1:** Reaction rates used to generate synthetic trajectories.

| Rate | Value |
| --- | --- |
| $g$ ($\text{s}^{-1}$) | $5.0 \times 10^{-2}$ |
| $r$ ($\text{s}^{-1}$) | $3.0 \times 10^{-3}$ |
| $f$ ($\text{s}^{-1}$) | $5.0 \times 10^{-1}$ |
| $b$ ($\text{s}^{-1}$) | $10^{-2}$ |
| $\Delta t$ (s) | 60 |
| $T$ (days) | 1 |

### MaxCal Model Descriptions

#### Minimal model 1 (M1)

Our minimal Maximum Caliber (MaxCal) model is built on three basic features within the system: 1) protein production, 2) protein degradation, and 3) cyclic repression (see Figure 1). To account for protein production, each protein species has its own production-path variable, *ℓ_α_*, *ℓ_β_*, and *ℓ_γ_*, that represents the number of proteins that are created in a discrete time interval of Δ*t* and ranges between zero and some predefined maximum, *M*. As for protein degradation, each species also has its own degradation-path variable, *ℓ_A_*, *ℓ_B_*, and *ℓ_C_*, that represents the number of preexisting proteins that remain after a discrete time interval of Δ*t* and ranges between zero and the initial number of proteins present, *N_A_*, *N_B_*, and *N_C_*. To constrain the behavior of the system, we introduce Lagrange multipliers to restrict the average values of each path variable, specifically *h_α_*, *h_β_*, *h_γ_*, *h_A_*, *h_B_*, and *h_C_*. All three proteins in equation 1 have symmetric reaction rates, hence we assume that all Lagrange multipliers are also symmetric across protein species (i.e. *h_α_* = *h_β_* = *h_γ_*, *h_A_* = *h_B_* = *h_C_*). Within the simplest model (M1), we use an additional Lagrange multiplier, *K_Aβ_*, to impose a correlation between the presence of one protein and the production of the next (clockwise direction in Figure1), specifically restricting the average values of *ℓ_β_ℓ_A_*, *ℓ_γ_ℓ_B_*, and *ℓ_α_ℓ_C_*. With these definitions, the path entropy or “caliber” is given by:

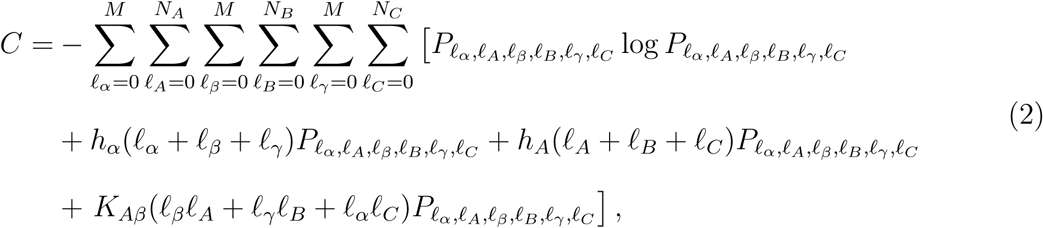

where *P_ℓα_*, *ℓ_A_*, *ℓ_β_*, *ℓ_B_*, *ℓ_γ_*, *ℓ_C_* is the probability of observing a path with a particular combination of *ℓ_α_*, *ℓ_A_*, *ℓ_β_*, *ℓ_B_*, *ℓ_γ_*, and *ℓ_C_*. Maximizing this caliber function with respect to the path probability distribution yields

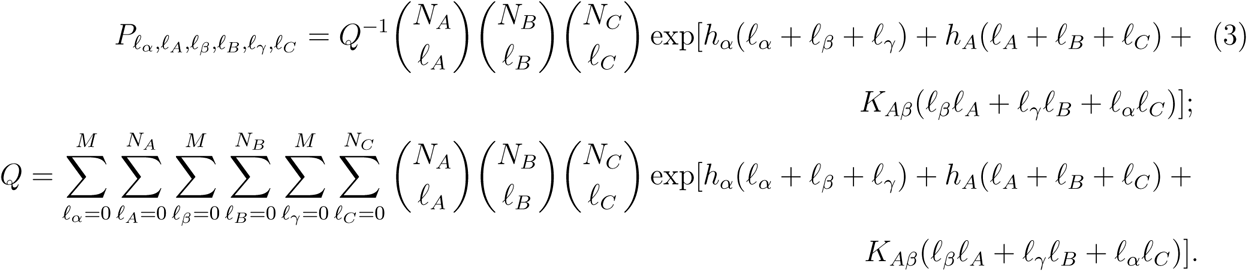

Stochastic protein expression trajectories similar to experiment/simulation (Figure 2) are generated from the path probability distribution by randomly selecting a path according to this distribution, creating and destroying the number of proteins corresponding to that path, and advancing the time of the system by Δ*t*. These path probabilities over the discrete time interval of Δ*t* can also be propagated in time to calculate the likelihood of stochastic trajectories (from experiment or simulation) using a discretized form of Finite State Projection (FSP).^54^ See Supporting Information for details on this formalism of FSP.

**Figure 2:**
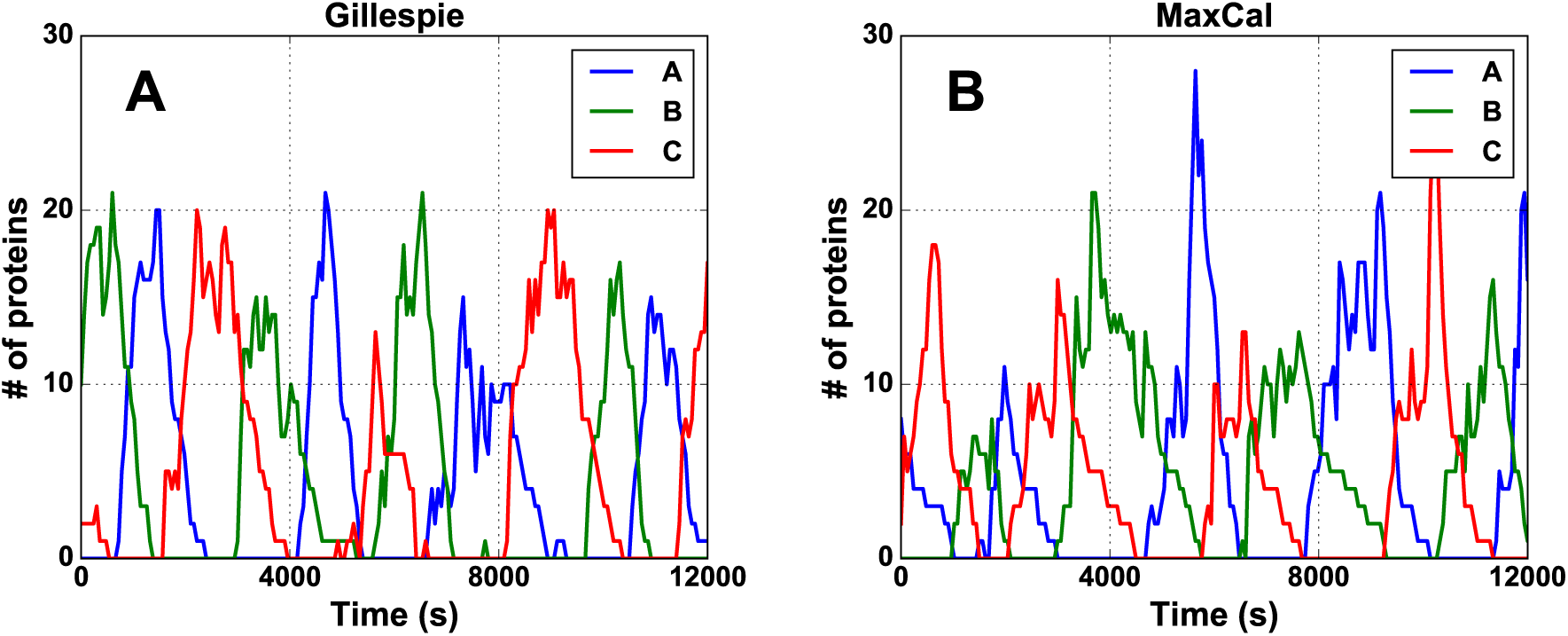
Sample Traces of MaxCal agre1e well with simulation. Direct comparison of synthetic input trajectories generated from a known model using Gillespie algorithm (**A**) and simulated MaxCal trajectories (**B**). Representative MaxCal parameters: *h_α_* = *−*0.460, *h_A_* = 1.812, *K_Aβ_* = *−*4.635, *M* = 17.

The utility of FSP is particularly advantageous in selecting values of *h_α_*, *h_A_*, *K_Aβ_*, and *M* via maximum likelihood (ML). Given a stochastic trajectory of protein expression (via experiment or simulation), how does one select the optimum values for these parameters that best represent that trajectory? Segmenting input trajectories frame-by-frame, equation 3 can be used to find the probability of each frame occurring randomly, and from there, the likelihood (*℡*) of the entire trajectory can be constructed for a particular set of values of these parameters. The parameter set that maximizes the likelihood of the observed trajectory is selected as the representative set. However, experiment and synthetic trajectories generated using Gillespie algorithm do not have upper limits on protein production analogous to *M* in MaxCal. Rare jumps in protein number over a single time step of Δ*t* (i.e. greater than *M*) disproportionately punish the likelihood of otherwise suitable parameters. To account for this, we use FSP (see Supporting Information) to calculate transition probabilities (*P_j→i_*_,*m*_) over multiple time increments (*m* frames) as

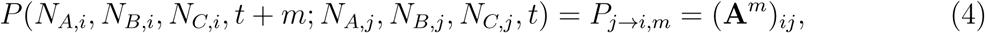

where **A** is the state reaction matrix described in the Supporting Information. This will allow MaxCal to realize large jumps in protein numbers over multiple frames and avoid erroneous penalization of Lagrange multipliers due to such rare events. Consequently, we calculate the likelihood *ℒ* of observing the experimental/simulated trajectory in increments of *m* frames as

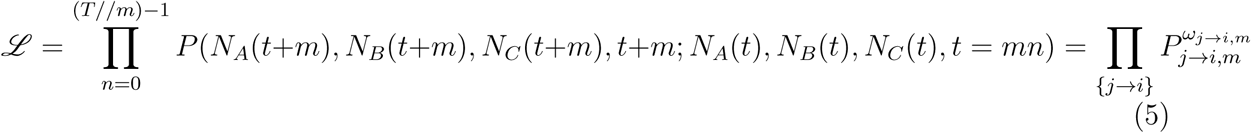

where *N_A_*(*t*), *N_B_*(*t*), and *N_C_* (*t*) are the number of *A*, *B*, and *C* proteins respectively at frame *t*, // is the standard floor division operation, *ω_j→i,m_* is the total number of *j → i* state transitions over *m* frames, and the second product is over all possible state transitions. Representative values of *h_α_*, *h_A_*, *K_Aβ_*, and *M* are selected by maximizing this likelihood with respect to these parameters. In order to capture the proper phase shift, i.e. oscillatory ordering between *A*, *B*, and *C*, maximum of *m* should be approximately the number of frames corresponding to one third of the average oscillatory period.

#### Model 2 (M2)

The description above illustrates application of the simplest MaxCal model, but models with higher complexity accounting for different crosstalk among *A*, *B*, and *C* are also possible. MaxCal formalism can be used to systematically incorporate these in the model by modifying equations 2 and 3. In the absence of direct knowledge about the underlying circuit, as would be the case with experimental data, it is not possible to exclude the possibility that each gene has some degree of auto-feedback in addition to the repression introduced in M1. To account for this possibility, we now present a second model (M2) that enforces a correlation between the presence of each protein species (*ℓ_A_*, *ℓ_B_*, *ℓ_C_*) and its own production (*ℓ_α_*, *ℓ_β_*, *ℓ_γ_*). This constraint is imposed by an additional Lagrange multiplier, *K_Aα_*, modifying the caliber function in equation 2 to the one below:

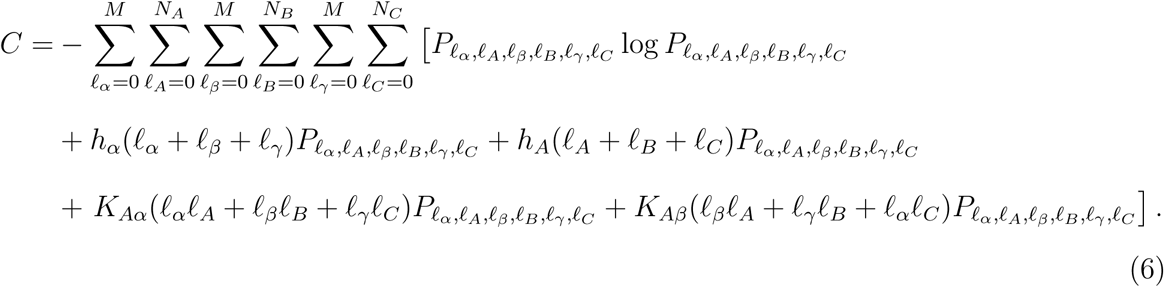

The corresponding caliber-maximized path probability function becomes

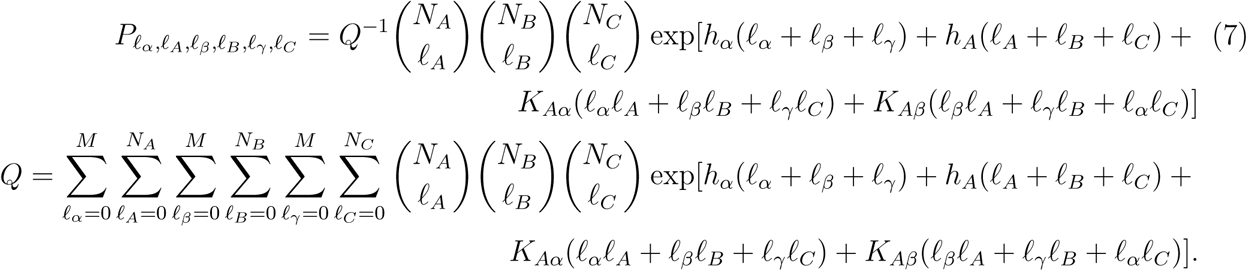

With this modified path probability distribution, FSP and ML formalism are executed as above.

#### Model 3 (M3)

Next, we present MaxCal model 3 (M3) by incorporating circular promotion/repression in the opposite direction of the preexisting oscillatory order (*A → B → C*). This is done by introducing additional symmetric couplings: the presence of protein A with the production of protein C, the presence of protein C with the production of protein B, and the presence of protein B with the production of protein A. Specifically, we constrain the average of *ℓ_A_ℓ_γ_*, *ℓ_C_ℓ_β_*, and *ℓ_B_ℓ_α_* imposed by the Lagrange multiplier *K_Aγ_*, with the new caliber function defined as

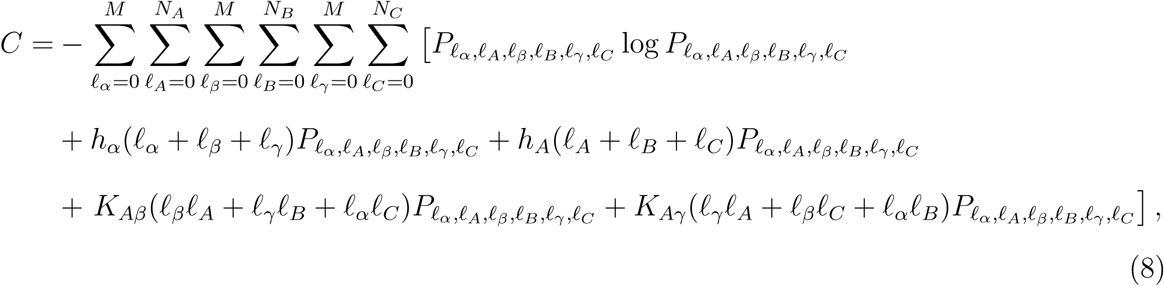

The caliber-maximized path probability is given by

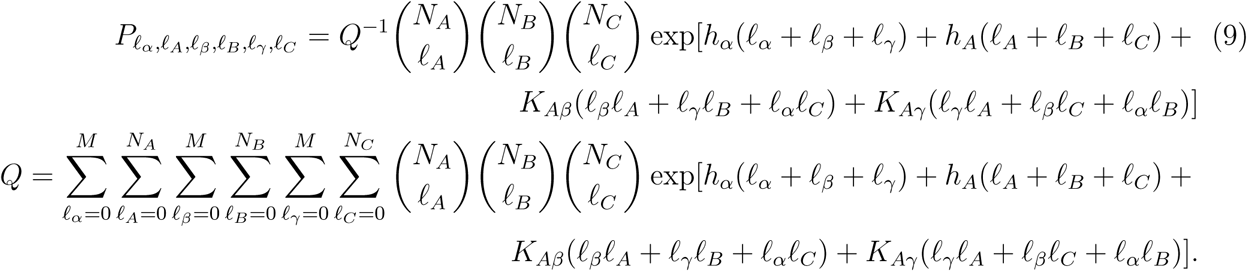

Again, FSP and ML are executed in the same way as before using the new set of parameters.

#### Model 4 (M4)

Finally, we present model 4 (M4) that accounts for all sets of crosstalk between existing protein numbers and production variables up to the second order. Thus, M4 allows all possible coupling and avoids ad-hoc guesswork of M1, M2, and M3 when underlying details are not known a priori. The caliber function for the most general model M4 is given by

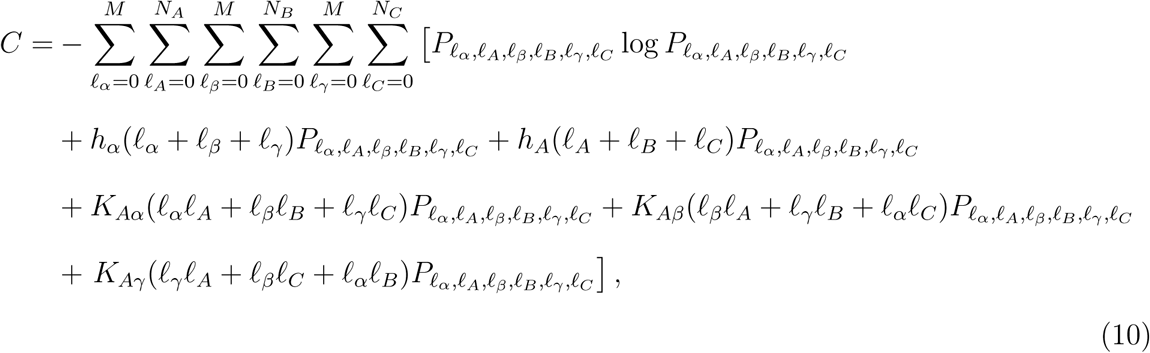

with the caliber-maximized path probability function given as

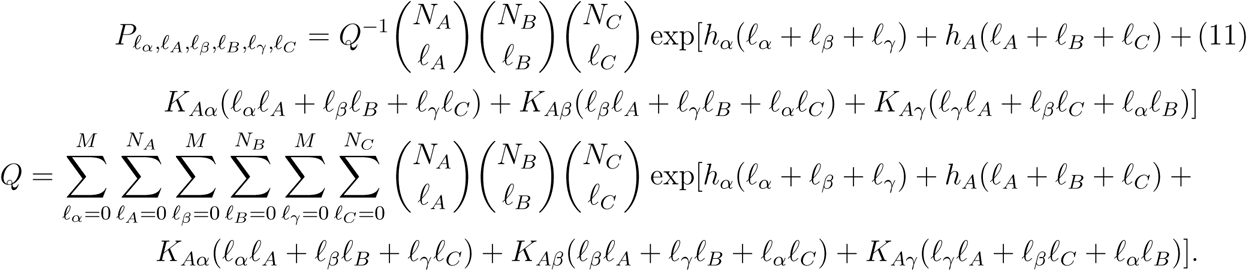

As before, when analyzing an observed trajectory, FSP and ML are carried out in the same way, but with respect to the six parameters *h_α_*, *h_A_*, *K_Aα_*, *K_Aβ_*, *K_Aγ_*, and *M*.

### Predictions and metrics used to assess MaxCal models

#### Inferring effective rates

To benchmark predictive powers of different MaxCal models, we consider effective protein production and degradation rates and compare them with the known values used to generate the synthetic data. Given specific protein numbers for each protein type (*A*, *B*, *C*), the transition probabilities described in equation 3 allow us to calculate effective protein production and degradation rates for each species. In the case of production of *A*, such a rate relates to the average of *ℓ_α_* as

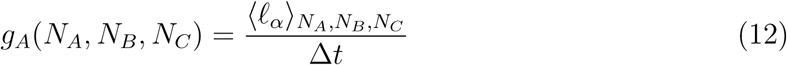

with symmetric equations for *B* and *C*, where 〈…〉_i,j,k_ represents the average of a quantity given that there are *i*, *j*, and *k* numbers of *A*, *B*, and *C* proteins initially present respectively. Next, we define the effective protein production rate (*g*_eff_) to compare with *g* in equation 1. Specifically, we recognize that a protein species’ basal production should be optimally observed when no other protein species are present to influence its production.

Thus, we define

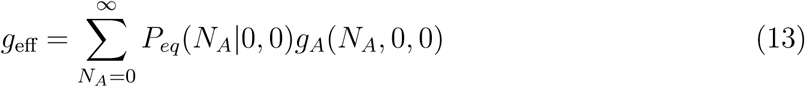

with symmetric equations for *B* and *C*, where *P_eq_*(*i|*0, 0) is the probability at relative equilibrium (calculated via discrete FSP) that there exists *i* number of proteins of any one of the three species given that the other two protein species are not present in the system. A similar approach can be taken to calculate an effective degradation rate for *A*, specifically relating to the average of *ℓ_A_* as

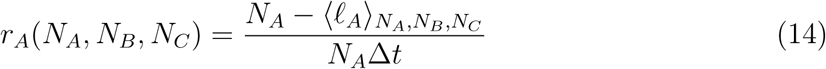

with symmetric equations for *B* and *C*. Since proteins should degrade at generally the same rate no matter what protein concentrations are present, we can predict an effective protein degradation rate (*r*_eff_) to compare with *r* in equation 1 by taking a weighted average of these values over the entire phase space of *N_A_*, *N_B_*, and *N_C_*. Thus, we define

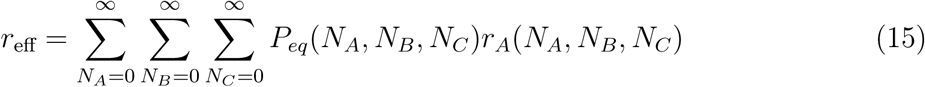

with symmetric equations for *B* and *C*, where *P_eq_*(*i*, *j*, *k*) denotes the probability at relative equilibrium (after transient dynamics) of having *i*, *j*, and *k* numbers of *A*, *B*, and *C* proteins respectively.

#### Inferring effective feedback

One novel aspect of the MaxCal formalism is the ability to calculate effective feedback metrics for different interactions. For example, the relative amount of cyclic (*A → B → C*; clockwise in Figure 1) repression in the system is quantified by calculating the Pearson correlation coefficient between the presence of one protein (*ℓ_A_*) and the production of the next (*ℓ_β_*). Specifically, we define this metric as

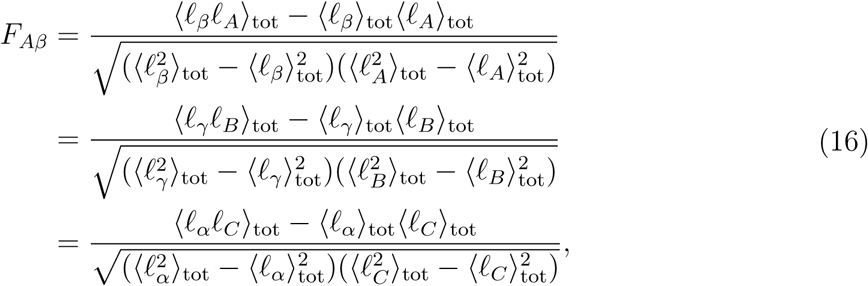

where

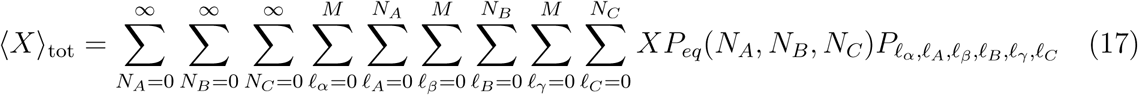

with *X* being any function of the stochastic variables of interest used to describe protein production and degradation. Furthermore, other forms of feedback can also be defined in a similar manner. For example, auto-feedback is estimated as

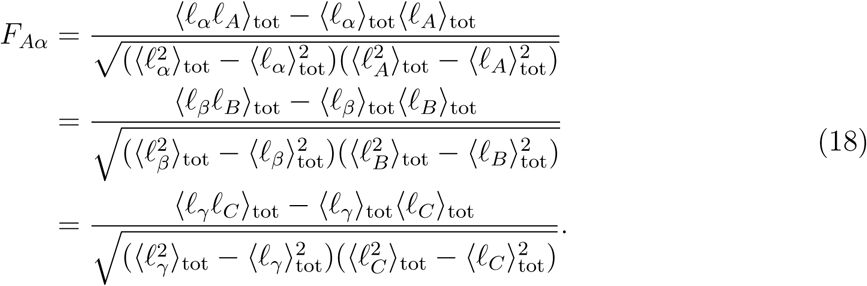

The cyclic promotion/repression in the anti-clockwise direction (*C → B → A*) is defined as,

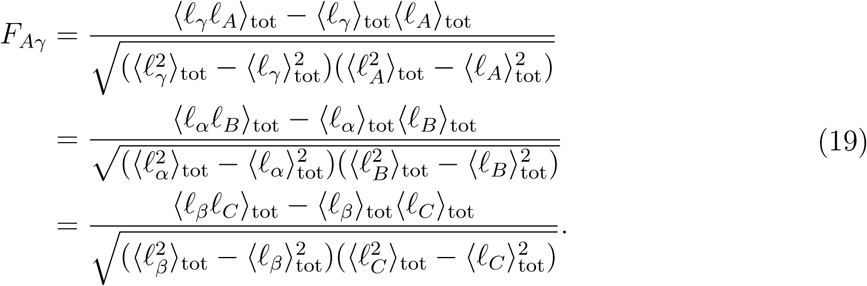

#### Trajectory entropy for assessment

For further quantitative comparison of how well MaxCal describes the observed features of the synthetic data, we consider several metrics in addition to the predictive and biophysically relevant metrics described above. Similar to our previous work,^34,35^ we compute two different path entropy metrics. The single-trajectory path entropy, *S*_1_, considers the protein number trajectories of *A*, *B*, and *C* independently of each other and is measured in bits as

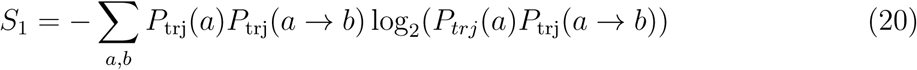

where *P*_trj_(*a*) is the probability of having *a* proteins (of one kind, *A* or *B* or *C*) at any point in any of the trajectories and *P*_trj_(*a → b*) is the probability of transitioning from *a* proteins to *b* in these trajectories. Next, we consider total path entropy, *S*_tot_, using the joint probability of all three of the protein number trajectories together. *S*_tot_ is calculated as

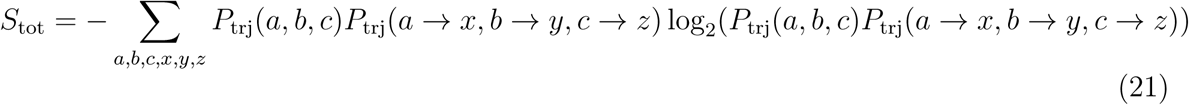

where *P*_trj_(*a*, *b*, *c*) is the joint probability of having *a*, *b*, and *c* numbers of *A*, *B*, and *C* proteins respectively during the course of the trajectories and *P*_trj_(*a → x*, *b → y*, *c → z*) is the probability of transitioning at those timepoints from that state to *x*, *y*, and *z* numbers of *A*, *B*, and *C* proteins respectively.

#### Comparing oscillatory frequency

Due to the oscillatory nature of these trajectories, we will also compare the oscillatory periods and peak heights between the input synthetic data and MaxCal inferred model. We switch from time domain to frequency domain by taking the Fast Fourier Transform (FFT) of the input time trace. From the frequency spectrum, we choose the frequency with the highest amplitude as the representative frequency (*f*_rep_) of the system. Consequently, the representative period is calculated as *τ*_rep_ = 1*/f*_rep_. Peak heights are measured by segmenting individual protein number trajectories into “peaks” and “valleys” by setting a threshold at 25% of the global maximum of protein number. Once a trajectory crosses this threshold, it is considered to be in a “peak” until it returns to zero, at which point it is considered to be in a “valley” until it crosses the threshold again. The maximum value of each peak section is then noted as the peak height.

#### Bayesian Information Criteria (BIC)

Different models have different numbers of parameters. For example, model 4 has six parameters while model 1 has four parameters. How do we justify addition of new parameters ? For a comparative analysis between models with different numbers of parameters, we provide two other metrics. First, we report the negative log likelihood of each model and the model with the highest likelihood is chosen. Next, we use Bayesian information criteria^55^ (BIC) to appropriately penalize models with more parameters even if they yield a higher likelihood. Specifically, BIC is defined as

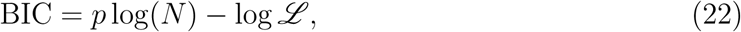

where *p* is the number of parameters, *N* is the number of measurements in the dataset, and *ℒ* is the likelihood. The first term in BIC penalizes the addition of each parameter while the second term rewards improved likelihood. Thus, BIC provides an objective way to select models with different parameters and different likelihoods.

### Analyzing typical data recorded in fluorescence

Models 1 (M1) through 4 (M4) assume that input trajectories are available in protein number. However, typical experiments record trajectories in fluorescence, not in protein numbers. Fluorescence per protein can be noisy and couples with gene expression noise. In order to apply MaxCal to infer underlying details of the circuits using gene expression noise, we must account for the fluorescence fluctuation in our ML parameter estimation protocol. We include the probability distribution of fluorescence collected per protein into our likelihood function to directly calculate the probability of observing a particular fluorescence trajectory rather than a particular protein number trajectory. Models 5 (M5) through 8 (M8) describe the equivalent of M1 through M4 but applied to fluorescence trajectories instead of protein number.

We test the inference capabilities of the fluorescence-based MaxCal models (M5, M6, M7, M8) using synthetic fluorescence trajectories mimicking real data. These synthetic trajectories were generated by purposefully “corrupting” the protein number trajectories (generated earlier from equation 1) with a typical distribution of fluorescence per protein. For simplicity, we assume this distribution to be Gaussian with an average of *f*_0_ and a standard deviation of *σ*. With this assumption, each time-point (with protein number *N_t_*) can be assigned a random fluorescence value based on convolution of *N_t_* individual fluorescence per protein distributions, i.e. another Gaussian distribution with an average of *N_t_f*_0_ and a standard deviation of 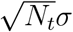. We consider relative noise levels (*σ/f*_0_) of 30% that are typical in experiments.^34^

To incorporate fluorescence into our likelihood calculation for inference, we combine path probabilities from MaxCal with the probability distribution of fluorescence per protein, similar to our earlier work.^34,35^ We assume that the average and standard deviation of the fluorescence distribution are known via low-intensity fluorescence experiments. ^56–68^ With this information, equation 5 is modified as,

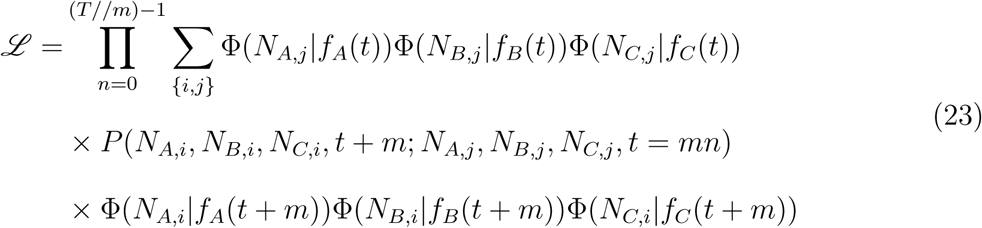

where *f_A_*(*t*), *f_B_*(*t*), and *f_C_* (*t*) are the fluorescences measured at time *t* from recorded trajectories for *A*, *B*, and *C* proteins respectively, and Φ(*N |f*) is the conditional probability that *N* proteins are present given that a fluorescence of *f* was observed. This conditional probability can be calculated using Bayes’ theorem as

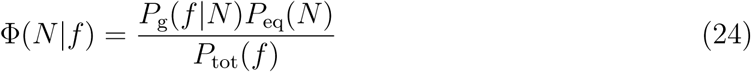

where *P*_g_(*f |N*) is the Gaussian fluorescence distribution with an average and standard deviation of *N f*_0_ and 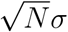, *P*_eq_(*N*) is the MaxCal-predicted protein number distribution at relative equilibrium (calculated via discretized FSP), and *P*_tot_(*f*) is the fluorescence probability distribution over the entire trace. Note that as before, *ℒ* is a function of the Lagrange multipliers and *M*, hence the optimal set of parameters for a given model (M5, M6, M7, or M8) is still obtained by maximizing the new *ℒ* function with respect to these parameters.

## Results and Discussion

### MaxCal models capture protein number trajectory fluctuations well and accurately infer the underlying model

The minimal model M1, with just the repression provided by *K_Aβ_*, reasonably reproduces features of the underlying trajectories. For example, M1 exhibits similar oscillatory frequencies (see Figure 3A), and oscillatory heights (see Figure 3B). It also recapitulates the dimensionally reduced protein number distribution for a given protein type (Figure 3C) and the joint probability of all three proteins (compare SI Figure 1-A and B).

**Figure 3:**
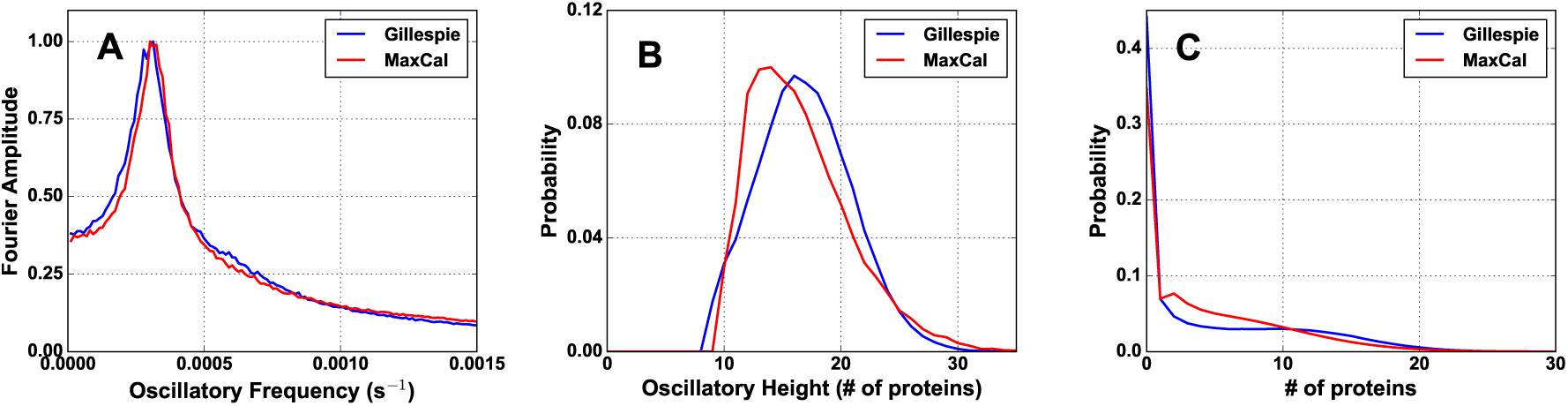
Model 1 agrees well with different characteristics of the synthetic data. Comparison between synthetic input trajectories generated from a known model using Gillespie algorithm (blue) and simulated MaxCal trajectories (red) of (**A**) oscillatory period, (**B**) oscillatory height, and (**C**) one-dimensional protein number distributions. Representative MaxCal parameters: *h_α_* = *−*0.459, *h_A_* = 1.810, *K_Aβ_* = *−*4.596, *M* = 16.

Table 2 further demonstrates the predictive power of MaxCal by comparing inferred degradation rates (*r*_eff_), production rates (*g*_eff_), representative periods (*τ*_rep_), average peak heights (〈*H*〉), and different entropy metrics (*S*_1_, *S*_tot_) against the “true” values. All the quantities, except *g*_eff_, are within 10% of the true value, highlighting the predictive power of M1. However, *g*_eff_ has an error of about 40%. The inferred feedback metrics (not readily available from the input model) exhibit the expected trend: strong negative feedback (*F_Aβ_*) between the presence of one protein and the production of the next (in the clock-wise direction in Figure 1), appreciable self-promotion (*F_Aα_*), and negligible feedback in the counter-clockwise direction of oscillation (*F_Aγ_*).

**Table 2:**
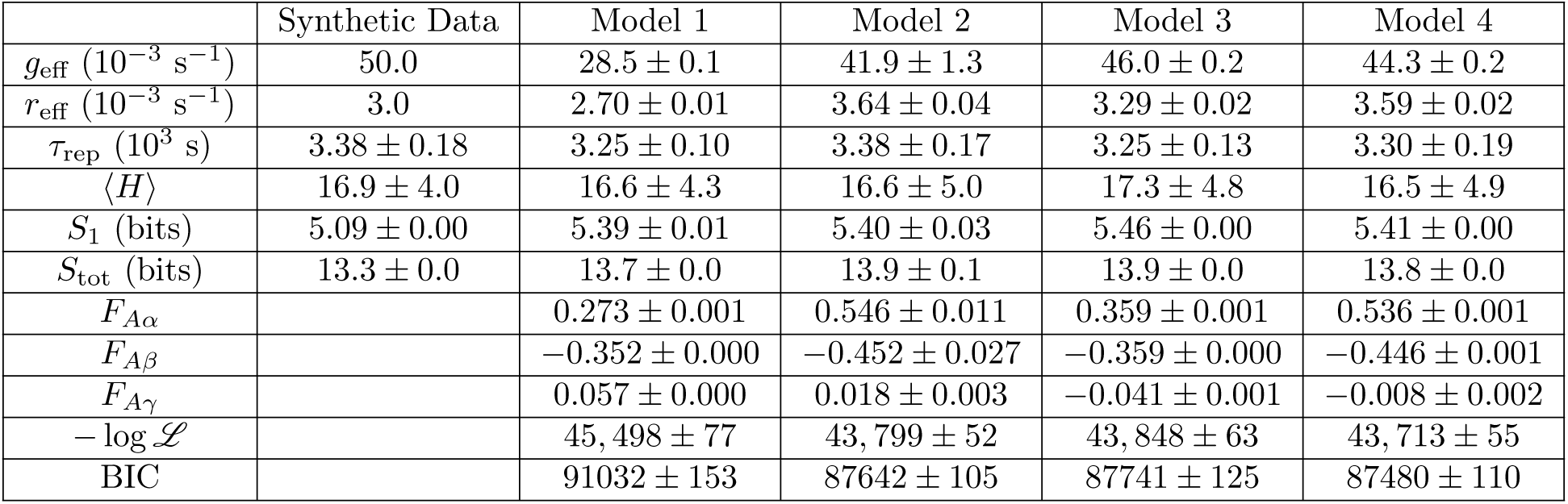
Comparison of “true” and MaxCal predicted rates and statistics from protein number based models. The last row gives the Bayesian Information Criteria to compare between models with different parameters.

Now we consider the performance of the higher order models, M2 and M3. The addition of *K_Aα_* and *K_Aγ_* in M2 and M3 respectively now show good agreement in all quantities. Interestingly, predicted *g*_eff_ agrees better with the “true” value, in contrast to M1. All feed-back metrics change slightly in magnitude, but the overall trends remain the same: *F_Aα_* is significantly positive, *F_Aβ_* is significantly negative, and *F_Aγ_* is negligible. Comparison of oscillatory frequencies (Figures 4A and 5A), oscillatory height distributions (Figure 4B, 5B), and protein number distributions (Figure 4C, 5C, SI Figure 1A, 1C, 1D) are also reasonable in reference to the input data. M2 and M3 (Figures 4A and 5A) tend to slightly overestimate the contributions of low frequencies, but the peak value dictating the representative frequency and period still coincides with that of the input data. The likelihood improves in both cases (smaller values of *−* log *ℒ* in Table 2 compared to M1). However, both M2 and M3 have an additional parameter compared to M1. To further evaluate the improvement of likelihood at the cost of an additional parameter, we calculate the BIC metric, and even with the BIC metric M2 and M3 are more likely, justifying the addition of the parameter. However, between M2 and M3, M2 is more likely. This is consistent with the fact that all three models (M1, M2, M3) predict *F_Aα_* to be higher in magnitude than *F_Aγ_* and suggests that *K_Aα_* (introduced in M2) is more relevant than *K_Aγ_* (in M3).

**Figure 4:**
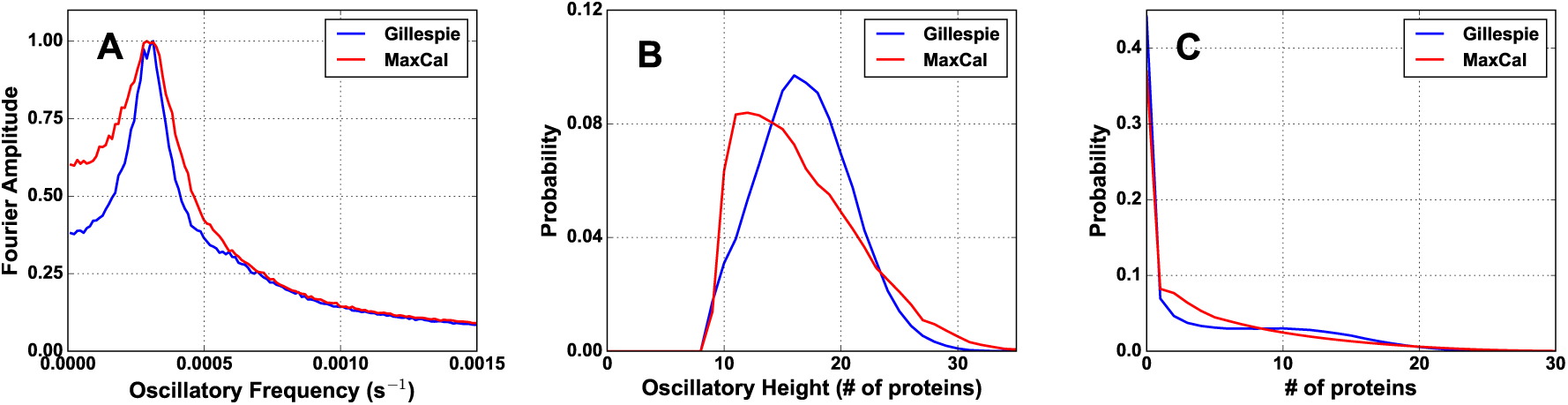
Model 2 agrees well with different characteristics of the synthetic data. Comparison between synthetic input trajectories generated from a known model using Gillespie algorithm (blue) and simulated MaxCal trajectories (red) of (**A**) oscillatory period, (**B**) oscillatory height, and (**C**) one-dimensional protein number distributions. Representative MaxCal parameters: *h_α_* = *−*0.389, *h_A_* = 1.533, *K_Aα_* = 0.109, *K_Aβ_* = *−*3.569, *M* = 3.

**Figure 5:**
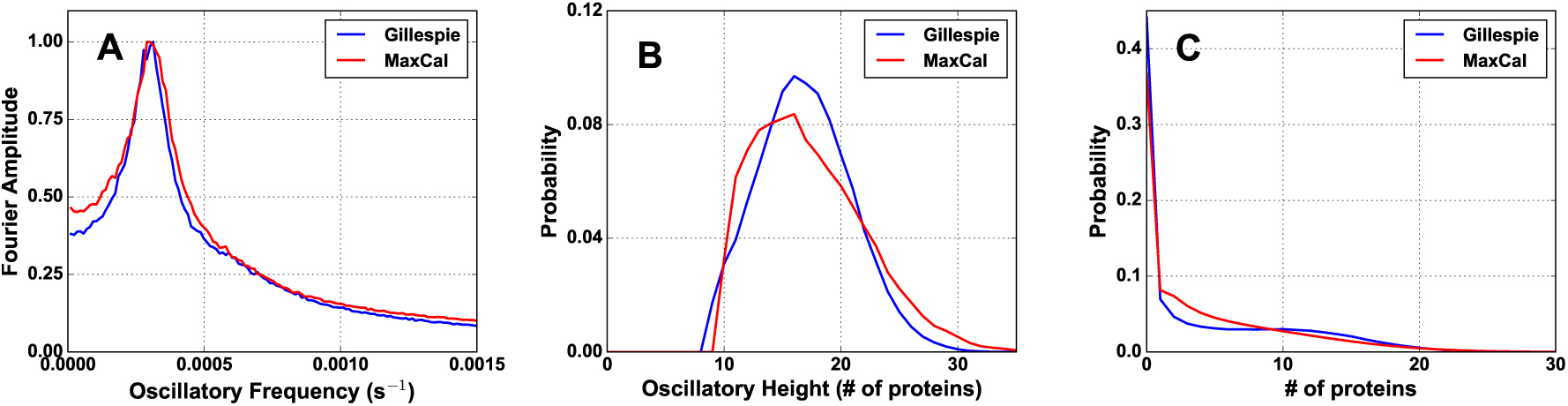
Model 3 agrees well with different characteristics of the synthetic data. Comparison between synthetic input trajectories generated from a known model using Gillespie algorithm (blue) and simulated MaxCal trajectories (red) of (**A**) oscillatory period, (**B**) oscillatory height, and (**C**) one-dimensional protein number distributions. Representative MaxCal parameters: *h_α_* = *−*0.230, *h_A_* = 1.655, *K_Aβ_* = *−*4.032, *K_Aγ_* = *−*0.043, *M* = 9.

Finally, M4 includes both *K_Aα_* and *K_Aγ_* in addition to *K_Aβ_* and provides a complete description of all possible forms of coupling within the system. Comparing the likelihood (second to last row in Table 2), we notice that M4 is the most likely of all models. However, M4 has an additional parameter compared to M2 and M3. Again, we use the BIC metric to assess whether it is justifiable to use the additional parameter in M4. Even with the BIC penalty, we notice that M4 is more likely than all other models. However, improvement beyond M2 and M3 in terms of distributions and statistics is minimal. All quantities in the last column of Table 2 remain within 20% of their true values. Protein number (Figure 6C, SI Figure 1E) and peak height distributions (Figure 6B) overlap well with that of the input trajectories. Lower frequencies are again more prominent in Figure 6A, while the representative frequency and period again compare well with the input. The values of feedback parameters follow the same trend as M1, M2, and M3.

**Figure 6:**
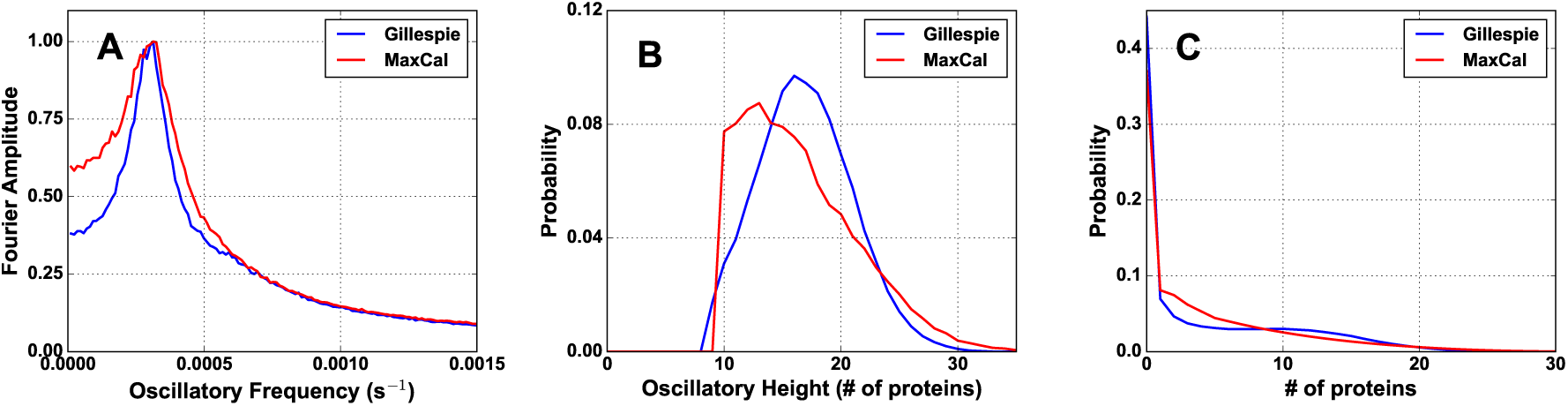
Model 4 agrees well with different characteristics of the synthetic data. Comparison between synthetic input trajectories (blue) and simulated MaxCal trajectories (red) of (**A**) oscillatory period, (**B**) oscillatory height, and (**C**) one-dimensional protein number distributions. Representative MaxCal parameters: *h_α_* = *−*0.389, *h_A_* = 1.537, *K_Aα_* = 0.063, *K_Aβ_* = *−*3.704, *K_Aγ_* = *−*0.014, *M* = 4.

We conclude that the minimal model M1 provides a reasonable description of the underlying system. However, M4 provides the best description of the system as judged by BIC and is preferred. M4 also includes all second-order crosstalk between production and protein number, providing a complete description of the system and avoiding the ad-hoc guesswork in M1, M2, and M3 which ignore different terms. If computational resources allow, M4 is more desirable, but the minimal M1 model can serve as a preliminary model when resources are limited.

### MaxCal models perform well even when data is in fluorescence

Results reported in the previous subsection used input trajectories provided in protein number, an unrealistic condition in experiment. To provide a more realistic assessment of MaxCal’s inferential capabilities, we generated fluorescence trajectories from the same exact protein number trajectories from the previous section by “corrupting” them with Gaussian fluorescence per protein distributions. These fluorescence traces were then provided to the MaxCal machinery as input and representative parameter values were selected using the ML formalism described in equation 23. Relative fluorescence noise levels were set to 30% to best match experimental conditions.

As seen in Figures 7-10, SI Figure 2, and Table 3, MaxCal still provides an accurate reproduction of the repressilator circuit even when gene expression noise is convoluted with fluctuations in fluorescence. Similar to M1, the minimal model M5 provides a reasonable representation of the underlying circuit. Models 6 through 8, representing the higher order MaxCal models applied to fluorescence trajectories, all show further improvement compared to M5. As before, we notice the introduction of the auto-feedback (*K_Aα_*) term in M6 increases the likelihood more compared to M7 (with the *K_Aγ_* term). While the choice of M6 over M7 is evident, we must revisit the choice of M8 over M6 due to parameter mismatch. We notice that the BIC metric for M6 and M8 are almost comparable, making selection between M6 and M8 ambiguous based on model performance. However, M8 is still desirable over M6 as it is the most general form and invokes no assumptions.

**Figure 7:**
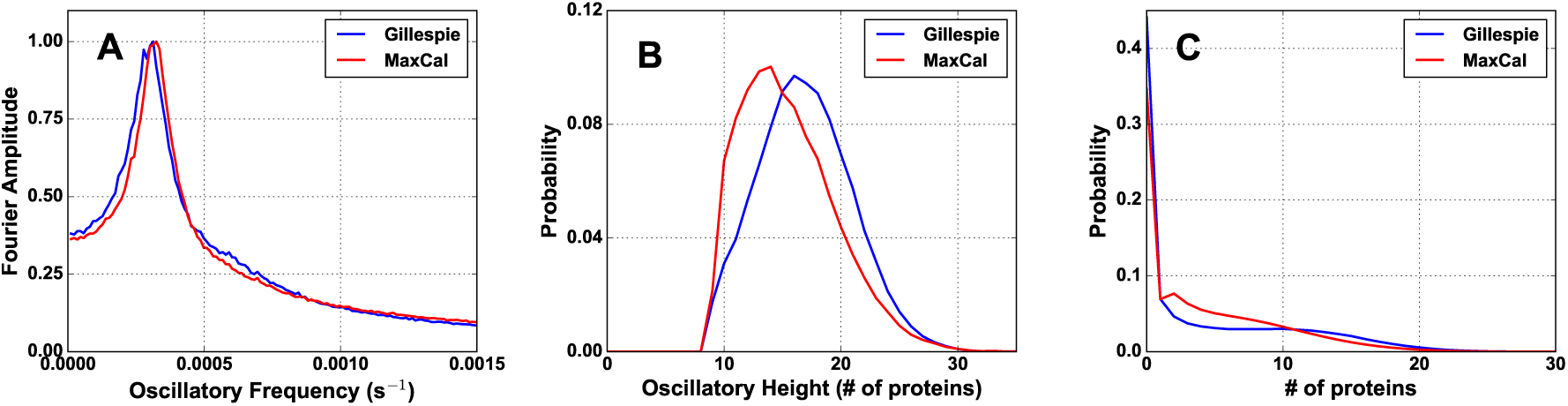
Model 5 agrees well with different characteristics of the synthetic data.. Comparison between synthetic input trajectories generated from a known model using Gillespie algorithm (blue) and simulated MaxCal trajectories (red) of (**A**) oscillatory period, (**B**) oscillatory height, and (**C**) one-dimensional protein number distributions. Representative MaxCal parameters: *h_α_* = *−*0.448, *h_A_* = 1.809, *K_Aβ_* = *−*4.646, *M* = 10.

**Figure 8:**
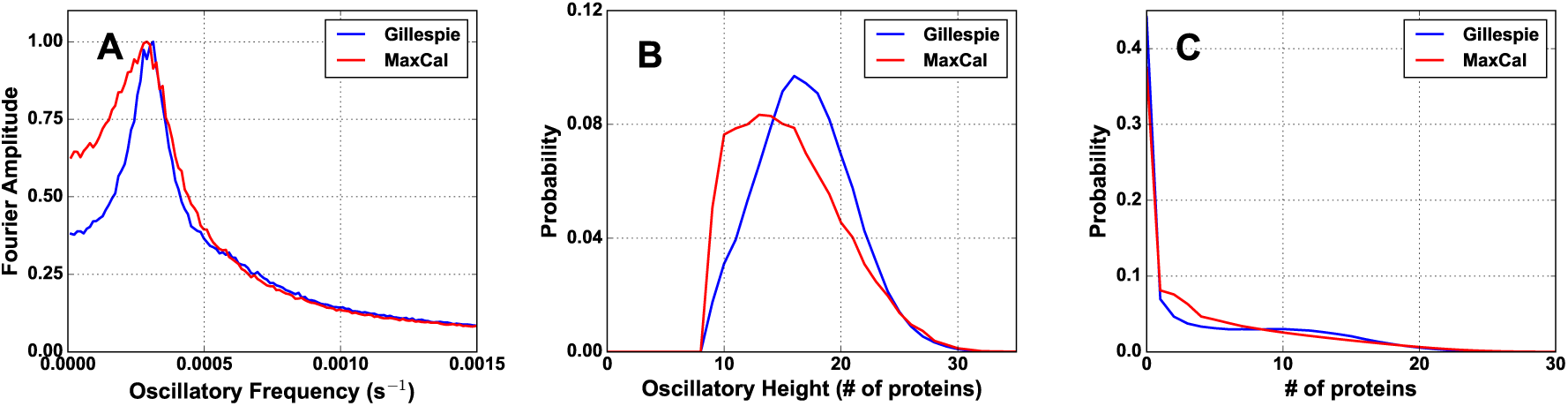
Model 6 agrees well with different characteristics of the synthetic data.. Comparison between synthetic input trajectories generated from a known model using Gillespie algorithm (blue) and simulated MaxCal trajectories (red) of (**A**) oscillatory period, (**B**) oscillatory height, and (**C**) one-dimensional protein number distributions. Representative MaxCal parameters: *h_α_* = *−*0.420, *h_A_* = 1.521, *K_Aα_* = 0.113, *K_Aβ_* = *−*3.780, *M* = 3.

**Figure 9:**
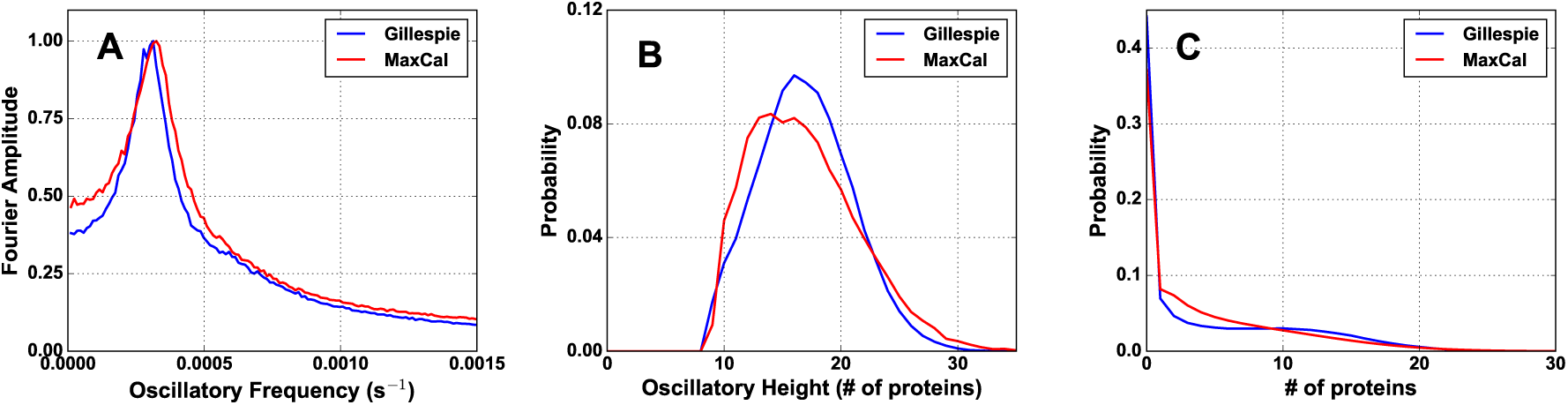
Model 7 agrees well with different characteristics of the synthetic data. Comparison between synthetic input trajectories generated from a known model using Gillespie algorithm (blue) and simulated MaxCal trajectories (red) of (**A**) oscillatory period, (**B**) oscillatory height, and (**C**) one-dimensional protein number distributions. Representative MaxCal parameters: *h_α_* = *−*0.193, *h_A_* = 1.623, *K_Aβ_* = *−*4.317, *K_Aγ_* = *−*0.048, *M* = 8.

**Figure 10.**
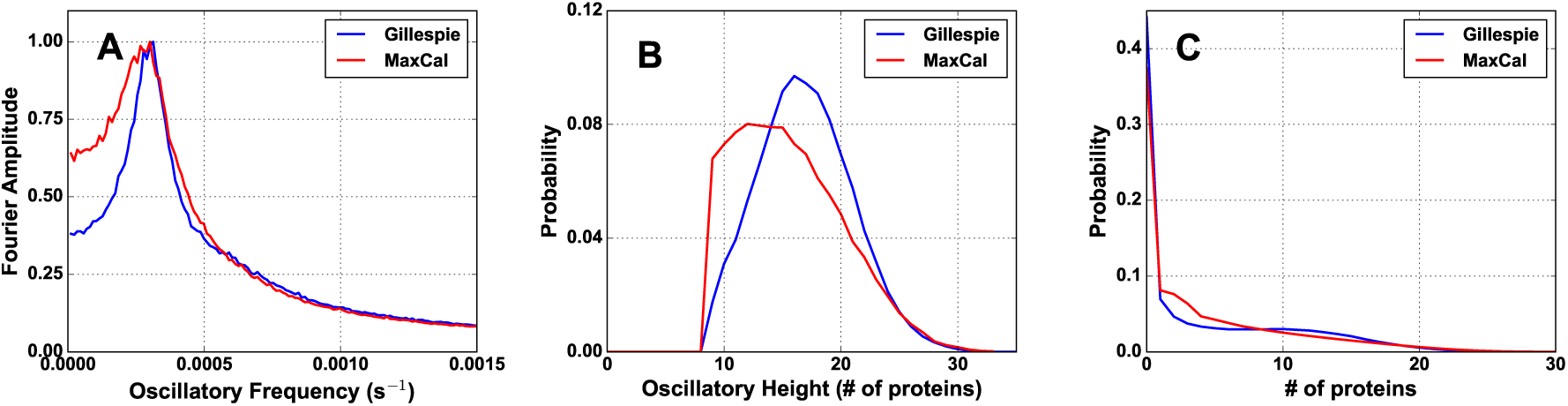
Model 8 agrees well with different characteristics of the synthetic data. Comparison between synthetic input trajectories generated from a known model using Gillespie algorithm (blue) and simulated MaxCal trajectories (red) of (**A**) oscillatory period, (**B**) oscillatory height, and (**C**) one-dimensional protein number distributions. Representative MaxCal parameters: *h_α_* = 0.456, *h_A_* = 1.510, *K_Aα_* = 0.116, *K_Aβ_* = 3.846, *K_Aγ_* = 0.004, *M* = 3.

**Table 3:** Comparison of “true” and MaxCal predicted rates and statistics from fluorescence based models. The last row reports Bayesian Information Criteria for a critical assessment of models with different parameters.

|  | Experiment | Model 5 | Model 6 | Model 7 | Model 8 |
| --- | --- | --- | --- | --- | --- |
| $g_{\text{eff}} (10^{-3} \text{ s}^{-1})$ | 50.0 | $28.1 \pm 0.0$ | $39.4 \pm 0.1$ | $46.7 \pm 0.3$ | $39.2 \pm 0.1$ |
| $r_{\text{eff}} (10^{-3} \text{ s}^{-1})$ | 3.0 | $2.70 \pm 0.00$ | $3.78 \pm 0.02$ | $3.45 \pm 0.02$ | $3.79 \pm 0.01$ |
| $\tau_{\text{rep}} (10^3 \text{ s})$ | $3.38 \pm 0.18$ | $3.17 \pm 0.09$ | $3.46 \pm 0.16$ | $3.08 \pm 0.07$ | $3.79 \pm 0.37$ |
| $\langle H \rangle$ | $16.9 \pm 4.0$ | $15.5 \pm 4.0$ | $15.7 \pm 4.5$ | $16.9 \pm 4.6$ | $15.6 \pm 4.6$ |
| $S_1$ (bits) | $5.09 \pm 0.00$ | $5.36 \pm 0.01$ | $5.33 \pm 0.00$ | $5.44 \pm 0.00$ | $5.34 \pm 0.01$ |
| $S_{\text{tot}}$ (bits) | $13.3 \pm 0.0$ | $13.6 \pm 0.0$ | $13.6 \pm 0.0$ | $13.8 \pm 0.0$ | $13.6 \pm 0.0$ |
| $F_{A\alpha}$ | | $0.285 \pm 0.000$ | $0.576 \pm 0.001$ | $0.374 \pm 0.001$ | $0.575 \pm 0.001$ |
| $F_{A\beta}$ | | $-0.369 \pm 0.000$ | $-0.498 \pm 0.001$ | $-0.370 \pm 0.000$ | $-0.497 \pm 0.001$ |
| $F_{A\gamma}$ | | $0.061 \pm 0.000$ | $0.018 \pm 0.001$ | $-0.047 \pm 0.001$ | $0.023 \pm 0.002$ |
| BIC | | $186980 \pm 225$ | $176830 \pm 318$ | $178082 \pm 312$ | $176824 \pm 315$ |

The success of MaxCal in building predictive models from noisy fluorescence data is non-trivial for three reasons. First, an increase in the number of species for which data is given in noisy fluorescence enhances the degree of obfuscation. Second, as seen in the previous subsection, the presence of three genes and their interdependence also requires using a joint distribution of protein numbers. Consequently, the coupling between these two sources of noise that individually scale with the number of species presents a formidable challenge to the inference machinery. By successfully decoupling all these different sources of fluctuations and building accurate models of the underlying network, we demonstrate scalability of MaxCal beyond the two-gene networks studied earlier.

## Conclusion

We have built multiple MaxCal models of three-gene feedback networks that exhibit oscillation in protein numbers. The simplest model (M1) uses only three constraints: protein production, degradation, and feedback (between the presence of *A* proteins and production of *B*; presence of *B* proteins and production of *C*; presence of *C* proteins and production of *A*). Additional coupling between different species were introduced in higher order models (M2, M3, and M4) with M4 allowing all possible forms of crosstalk between different species. Next, we tested inferential ability of these models by using a synthetic trajectory generated from a detailed reaction network model involving nine species (three promoters each in the active and inactive form and the three expressed proteins) with known parameters. This serves as a proxy for experimental data and is used for benchmarking MaxCal’s predictive power. By integrating MaxCal with maximum likelihood (ML) formalism, we show that MaxCal + ML can use the full information content of trajectory data and extract several important model parameters such as the effective degradation rate and production rate. All models – including the minimal model M1 – reasonably predict these effective rates when compared against the known values used to generate the synthetic data. MaxCal models also predict effective feedback parameters between different species, a feature not afforded to traditional modeling approaches. These predictive quantities demonstrate MaxCal’s ability to learn from the stochastic data about underlying details that are not directly available from data. Finally, we notice that MaxCal models can capture the underlying fluctuations of the stochastic trajectory and consequently predict several distributions of physical observables such as protein number distribution, peak height distribution, and frequency spectrum, that agree well with the input data. However, M4 most accurately captures fluctuation in the data as assessed by the Bayesian Information Criteria. Moreover, M4 avoids the ad-hoc guesswork inherent in M1, M2, and M3 that requires assuming that certain feedbacks are absent without any justification. Prediction of distributions as a function of model parameters further highlights MaxCal’s ability to tune circuits and optimize certain features, relevant for circuit design.

Next, we extend the applicability of our model to analyze experimental data, where stochastic trajectories are typically recorded in fluorescence and not in protein number. We do this by including a distribution for fluorescence per protein in the likelihood function of the previous models (M1 to M4). The modified models (M5 to M8) reasonably predict underlying parameters and fluctuations in data, highlighting MaxCal’s ability to deconvolute different sources of fluctuations. In summary, MaxCal provides the necessary framework to directly analyze raw experimental trajectories (in noisy fluorescence) without any data reduction and to build predictive models of underlying genetic networks involving up to three different genes. This opens up room for future applications modeling other network motifs – beyond switches and oscillators – with multiple genes interacting via complex feedback networks.

## Associated Content

### Supporting Information

The supporting information contains a discussion of a discrete-time FSP formalism and two graphs (Figure 1 and Figure 2). The two graphs show a comparison of the joint probability distribution of three proteins (*A*, *B*, and *C*) between the input data and inferred distribution using different MaxCal models. The probability distribution is shown as a three-dimensional heat map with dark red denoting higher probability. The first graph (Figure 1) is when the input data is given in particle number and the second graph (Figure 2) is when the input data is given in fluorescence.

## Acknowledgement

We acknowledge support from NSF (award number 1149992), RCSA (as a Cottrell Scholar), and the PROF grant from DU. Access to the High Performance Computing facility at DU is greatly appreciated, as well as the help of Ben Fotovich for GPU usage.

